## Supplemental information for "Physiological roles of endocytosis and presynaptic scaffold in vesicle replenishment at fast and slow central synapses"

**This file includes:**

**Supplementary Data**

- Figure S1 to S6
- Table S1 to S2

**Supplementary Text**

- Resources table

**Supplementary figure and table details:**

**Figure S1** (related to Figure 1). Effect of intra-terminal loading of Dynamin-1 PRD peptide on endocytosis at the calyx of Held

**Figure S2** (related to Figures 2, 4, and 6). Effect of endocytic inhibitors or scaffold cascade inhibitors on basal EPSC amplitude at the calyx and hippocampal CA1 synapses

**Figure S3** (related to Figure 2). Postsynaptic AMPA receptor saturation does not contribute to enhancement of synaptic depression by endocytic blockers at the calyx of Held

**Figure S4** (related to Figure 3). Endocytic blockers enhance STD but do not alter the recovery from STD time-course in 2.0 mM [$Ca^{2+}$] irrespective of PT or RT at the calyx of Held

**Figure S5** (related to Figure 3). Effects of Latrunculin application protocols and [$Ca^{2+}$] on STD

**Figure S6** (related to Figure 6). Vacuolar ATPase blockers, Bafilomycin or Folimycin does not affect the short-term synaptic facilitation at the hippocampal CA1 synapse

**Table S1** (related to Figures 1, 3 and S1). Membrane capacitance and calcium current amplitudes with or without endocytic blockers or presynaptic scaffold inhibitors during slow, fast-accelerating, or fast endocytosis

**Table S2** (related to Figures 2 and 4). Parameters of recovery from STD at 37°C and 1.3 mM $Ca^{2+}$ with or without endocytic- or presynaptic scaffold-blockers

**Figure S1. Effect of intra-terminal loading of Dynamin-1 PRD peptide on endocytosis at the calyx of Held**

**(A)** Schematic illustration of presynaptic membrane capacitance ($C_m$) recording and loading of dynamin-1 proline-rich domain (Dyn1-PRD) peptide (sequence: PQVPSRPNRAP) into a calyceal terminal (maroon) through a recording pipette.

**(B)** Average traces of slow endocytic membrane capacitance changes ($\Delta C_m$) in response to a 5-ms depolarizing pulse (stepping from -70 mV to +10 mV), without (control, black) or with dynamin-1 PRD peptide (1 mM, waiting 4-5 min after whole-cell rupture, maroon) at the calyx of Held presynaptic terminal in slices from P13-15 mice at physiological temperature (PT, 37°C) and in 2.0 mM $Ca^{2+}$ aCSF. Bar graphs on 3rd and 4th panels from left show magnitude of exocytic capacitance change ($\Delta C_m$) and calcium current in the presence of Dyn1-PRD peptide showing no difference from control (Table S1). Rightmost bar graph indicates that endocytic decay rate in dynamin PRD peptide (2.3 ± 2.2 fF; n = 5; $p < 0.001$, Student's t-test), was significantly slowed compared to control (28.8 ± 3.7 fF; n = 5).

**(C)** Average fast-accelerating endocytosis traces elicited by a train of 20-ms pulses (x15) at 1 Hz in control and in the presence of dynamin PRD peptide. In the 3rd panel, mean endocytic rate (fF/s) calculated from the slope of $C_m$ decay 0.45-0.95 ms after each stimulation pulse in control and under Dyn-1 PRD peptide are superimposed. The bar graph in the 4th panel shows the average endocytic decay rate (between #12-15, bar in 3rd panel from left) that was significantly less in Dyn1-PRD peptide (132 ± 10 fF; n = 7; $p = 0.001$) than control (251 ± 28.3 fF; n = 6).

**(D)** Averaged traces of fast endocytosis induced by a 10-Hz train of 20-ms pulses (x10) in control and in the presence of Dyn1-PRD peptide. In the 3rd panel, superimposed $C_m$ traces are shown in expanded time scale. The cumulative exocytic change ($\Delta C_m$) in the presence of Dyn1-PRD peptide was significantly lower than control (Table S1). Rightmost bar graph indicates that the fast endocytic decay rate was significantly slowed in the presence of Dyn1-PRD peptide (161 ± 27.3 fF; n = 6; $p = 0.003$, t-test) compared to control (346 ± 40.1 fF; n = 6).

**Figure S2. Effect of endocytic inhibitors or scaffold inhibitors on basal EPSC amplitude at the calyx and hippocampal CA1 synapses.**

**(A)** Basal mean amplitude of EPSCs recorded from the calyx of Held in control (6.6 ± 0.54 nA, n = 22) was unaffected by Dynasore (7.6 ± 0.67 nA; n = 12; $p = 0.24$),

Pitstop-2 (6.6 ± 0.61 nA; n = 8; $p = 0.96$), Latrunculin-B (6.4 ± 0.67 nA; n = 8; $p = 0.8$), or ML141 (6.7 ± 0.64 nA; n = 12; $p = 0.9$).

**(B)** Basal mean amplitude of EPSCs recorded from hippocampal CA1 area in control (78 ± 9.5 pA; n = 17) was increased by Dynasore (165 ± 12 pA; n = 10; $p < 0.001$, Student's t-test), but not by Pitstop-2 (70 ± 10 pA; n = 14; $p = 0.6$), Lat-B (85 ± 12 pA; n = 12; $p = 0.63$), or ML141 (76 ± 10 pA; n = 10; $p = 0.88$).

The statistical significance of all data in this figure was evaluated by one-way ANOVA and Student's t-test, with Bonferroni-Holm correction.

**Figure S3. Postsynaptic AMPA receptor saturation does not contribute to enhancement of synaptic depression by endocytic blockers at the calyx of Held**

The low affinity glutamate receptor ligand kynurenic acid (Kyn, 1 mM) attenuated the EPSC amplitude (left top panel). In the presence of Kyn, the magnitude of enhancement of synaptic depression by endocytic blockers was essentially the same as that in their absence (Figure 2).

**(A)** At 10-Hz stimulation, the steady state depression (stimulation #26-30) in the presence of Kyn or control (0.47 ± 0.02; n = 6) was slightly enhanced by Kyn + Dynasore (0.37 ± 0.02; n = 6; $p = 0.004$, Student's t-test), but not by Kyn + Pitstop-2 (0.41 ± 0.014; n = 7; $p = 0.052$).

**(B)** At 100-Hz stimulation, starting from the $2^{nd}$ stimulation (10 ms from onset) synaptic depression in control (0.97 ± 0.1, n = 5) was significantly enhanced in the presence of Dynasore (0.69 ± 0.02, n =6; $p = 0.012$) or Pitstop-2 (0.64 ± 0.056, n = 7; $p = 0.01$). The steady state depression (#26-30) in control (0.32 ± 0.02; n = 5) was significantly enhanced in the presence of Dynasore (0.19 ± 0.2; $p < 0.001$) or Pitstop-2 (0.18 ± 0.01; $p < 0.001$). One-way ANOVA and t-test, with Bonferroni-Holm correction was used to evaluate the statistical significance of all data in this figure.

**Figure S4. Endocytic blockers enhance STD but do not alter the recovery from STD time-course in 2.0 mM [$Ca^{2+}$] irrespective of PT or RT at the calyx of Held**

Effects of Dynasore and Pitstop-2 on short-term synaptic depression (STD; A1, B1, and C1) and on the time course of normalized recovery from STD (A2, B2, and C2) during 100-Hz (A, B) or 200-Hz (C) stimulation at PT (A, C) or RT (B).

**(A)** At PT (37°C) and in 2.0 mM $Ca^{2+}$ aCSF, the magnitude of EPSCs at 100 Hz show enhanced steady state depression in the presence of either Dynasore (0.23 ± 0.02; n

= 9; $p$ = 0.003, Student's t-test) or Pitstop-2 (0.23 ± 0.02; n = 11; $p$ = 0.002), in comparison to control (0.32 ± 0.02; n = 16). However, the kinetics of normalized recovery from STD ($\tau_{fast}$ and $\tau_{slow}$) in the presence of Dynasore ($\tau_{fast}$: 0.094 ± 0.024 s, n = 4, $p$ = 0.2, t-test) and ($\tau_{slow}$: 2.2 ± 0.5 s, $p$ = 0.3) or Pitstop-2 ($\tau_{fast}$: 0.1 ± 0.02 s, n = 9, $p$ = 0.2) and ($\tau_{slow}$: 1.7 ± 0.2 s; $p$ = 0.97) were not different from control ($\tau_{fast}$: 0.06 ± 0.014 s, n = 5) and ($\tau_{slow:}$ 1.7 ± 0.2 s).

**(B)** In 2.0 mM $Ca^{2+}$ at room temperature (RT; 22-24°C), Dynasore (0.12 ± 0.02; n = 6; $p$ = 0.016) or Pitstop-2 (0.10 ± 0.015; n = 5; $p$ = 0.008) enhanced STD (100 Hz) from control (0.19 ± 0.02; n = 11). The kinetics of recovery from STD ($\tau_{fast}$ and $\tau_{slow}$) in the presence of Dynasore ($\tau_{fast}$: 0.13 ± 0.02 s; n = 6; $p$ = 0.4) and ($\tau_{slow}$: 3.8 ± 0.7 s; $p$ = 1) or Pitstop-2 ($\tau_{fast}$: 0.17 ± 0.06; n = 5; $p$ = 0.7) and ($\tau_{slow}$: 4.2 ± 0.8 s; $p$ = 0.7) were unchanged from control ($\tau_{fast}$: 0.22 ± 0.09 s; n = 7) and ($\tau_{slow}$: 3.8 ± 0.6).

**(C)** During 200 Hz stimulation at PT and in 2.0 mM $Ca^{2+}$ aCSF, steady state STD was significantly enhanced by Dynasore (0.16 ± 0.014; n = 6; $p$ = 0.001) or Pitstop-2 (0.15 ± 0.017 s; n = 7; $p$ < 0.001) compared to control (0.25 ± 0.015; n = 10). However, the normalized recovery kinetics in control ($\tau_{fast}$: 0.09 ± 0.03s; $\tau_{slow}$, 2.5 ± 0.41s, n =7) were not-changed by Dynasore ($\tau_{fast}$: 0.11 ± 0.05s, $p$ = 0.7; $\tau_{slow}$, 2.2 ± 0.5 s, $p$ = 0.7, n = 6) or Pitstop-2 ($\tau_{fast}$: 0.1 ± 0.03s, $p$ = 0.8; $\tau_{slow}$: 3.5 ± 0.9 s, $p$ = 0.32, n = 6).

**Figure S5. Effects of Latrunculin application protocols and [$Ca^{2+}$] on STD**

**(A)** Two types of Latrunculin-A application protocols; perfusion (A1) and pre-incubation (A2).

**(B)** In 2.0 mM [$Ca^{2+}$], normalized amplitudes of EPSCs (30x at 100 Hz) in the presence (green circles) or absence (control, black circles) of Latrunculin-A (20 µM) under perfusion (B1) or pre-incubation (B2) protocols.

**(C)** In 1.3 mM [$Ca^{2+}$], normalized amplitudes of EPSCs (30x at 100 Hz) in the presence of Latrunculin-A (20 µM, green) or Latrunculin-B (15 µM, red) applied through perfusion or in their absence (control, black circles).

**(D)** Bar graphs for the magnitude of synaptic depression at #26-30 during 100 Hz stimulation in control (black), Latrunculin A (green) and Latrunculin B (red). Data from perfusion experiments (left and middle panels) and from pre-incubation experiments (right panel) are compared. Left panel, in 1.3 mM [$Ca^{2+}$]; middle and right panels, in 2 mM [$Ca^{2+}$]. Left panel: in 1.3 mM [$Ca^{2+}$], perfusion of Lat-A (0.18 ± 0.009; n=5; $p$ <0.001) or Lat-B (0.18 ± 0.013; n = 8; $p$ <0.001) equipotently enhanced depression from control (0.42 ± 0.025; n = 11). Middle and right panel: in 2.0 mM [$Ca^{2+}$], perfusion of Lat-A (0.19 ± 0.013; n=5; $p$ <0.001) significantly enhanced depression from control (0.32 ± 0.02; n = 16), whereas Lat-A preincubation (0.27 ± 0.018; n=4; $p$ = 0.16) was not different from control (0.32 ± 0.02). The magnitude of depression by Latrunculin A in 2.0 mM [$Ca^{2+}$] was significantly greater in perfusion than preincubation ($p$ = 0.008). Statistical significance was evaluated using one-way ANOVA and Student's t-test, with Bonferroni-Holm method of $p$ level correction.

**Figure S6. Vacuolar (v-) ATPase blockers, Bafilomycin or Folimycin does not affect the short-term synaptic facilitation at the hippocampal CA1 synapse**

**(A, B)** EPSCs (30x) evoked in hippocampal CA1 pyramidal cells by Schaffer collateral stimulation at 10 Hz **(A)** or 25 Hz **(B)** in the absence (control, black) or presence of v-ATPase blockers Bafilomycin (5 μM, 10-60 min, orange) or Folimycin (67 nM, 10-60 min, violet) at PT (37°C) and in 1.3 mM $Ca^{2+}$ aCSF. Panels from left to right: sample traces; normalized average EPSCs plotted against stimulation number. The bar graph indicates the magnitude of EPSC facilitation between-stimulations #26-30.

**(A)** At 10 Hz stimulation, the magnitude of synaptic facilitation at #26-30 in control (1.85 ± 0.17, n = 17) was unaffected by Bafilomycin (2.15 ± 0.11; n = 7; $p$ = 0.3) or Folimycin (2.2 ± 0.23; n = 8; $p$ = 0.21). The basal EPSC amplitude was significantly reduced in the presence of Bafilomycin (32.5 ± 6.1 pA; $p$ = 0.006) or Folimycin (40.0 ± 8.0 pA; $p$ = 0.014) than control (78 ± 9.5 pA).

**(B)** Also at 25 Hz, synaptic facilitation in control (2.65 ± 0.3; n = 16) was unchanged by Bafilomycin (2.5 ± 0.25; n = 6; $p$ = 0.8) or Folimycin (2.7 ± 0.18; n = 7; $p$ = 0.9).

One way ANOVA and Student's t-test, with Bonferroni-Holm correction was used to evaluate the statistical significance.

**Table S1. Membrane capacitance and calcium current amplitudes with or without endocytic blockers or presynaptic scaffold inhibitors during slow, fast-accelerating, or fast endocytosis**

**Table S2. Parameters of recovery from STD values at 37°C and 1.3 mM $Ca^{2+}$ with or without endocytic- or presynaptic scaffold -blockers**

**Resources Table:**

| REAGENT or RESOURCE | SOURCE | IDENTIFIER |
|---|---|---|
| Key Reagents/Peptide: | | |
| Dynasore | Abcam | Cat# ab120192 |
| Pitstop-2 | Abcam | Cat# ab120687 |
| ML141 | Abcam | Cat# ab145603 |
| Latrunculin B | Abcam | Cat# ab144291 |
| Folimycin (or Concanamycin A) | Abcam | Cat# ab144227 |
| Bafilomycin A1 | Cayman Chemical | Cat# 11038 |
| Dynamin-1 PRD peptide | GenScript | N/A (Custom ordered) |
| Strychnine HCl | Sigma-Aldrich | Cat# S8753 |
| Bicuculline Methiodide | Sigma-Aldrich | Cat# 14343 |
| D-AP5 | Tocris | Cat# 0106 |
| QX-314 bromide | Alomone Labs | Cat# Q-100 |
| DMSO, Sterile Filtered | Santa Cruz Chemicals | Cat# sc-359032 |
| Experimental Organism: | | |
| Mouse C57BL/6J | CLEA Japan | Cat# C57BL/6JJcl |
| Equipment's: | | |
| EPC 10 USB Patch Clamp Amplifier | Heka Elektronik | N/A |
| BX51WI upright microscope | Olympus | Cat# BX51WI |
| VT1200S Vibratome | Leica | N/A |
| Model 2100 Isolated Pulse Stimulator | A-M Systems | N/A |
| TC-344C Dual Channel Temp Controller | Warner Instruments | TC-344C |
| PatchStar Micromanipulator | Scientifica | PatchStar |
| Axiocam 506 mono camera system | Zeiss | Axiocam 506 mono |
| PIP 6 pipette puller | HEKA | PIP 6 |
| Borosilicate glass capillary (2.0 mm OD) | King Precision Glass Inc. | N/A |
| Software: | | |
| Patchmaster | HEKA | N/A |
| ZEN 2 Lite | Zeiss | N/A |
| IGOR Pro | Wavemetrics | N/A |
| Microsoft Excel | Microsoft | N/A |
| Corel Draw | Corel Corporation | N/A |

**Resource Availability:**

Further information about Resources/Reagents should be addressed to authors.

**Supplementary Information for**

### Physiological roles of endocytosis and presynaptic scaffold in vesicle replenishment at fast-signaling and slow-plastic synapses

Satyajit Mahapatra and Tomoyuki Takahashi

#### Supplementary Data

- Figure S1 to S6
- Table S1 to S2

### Figure S1

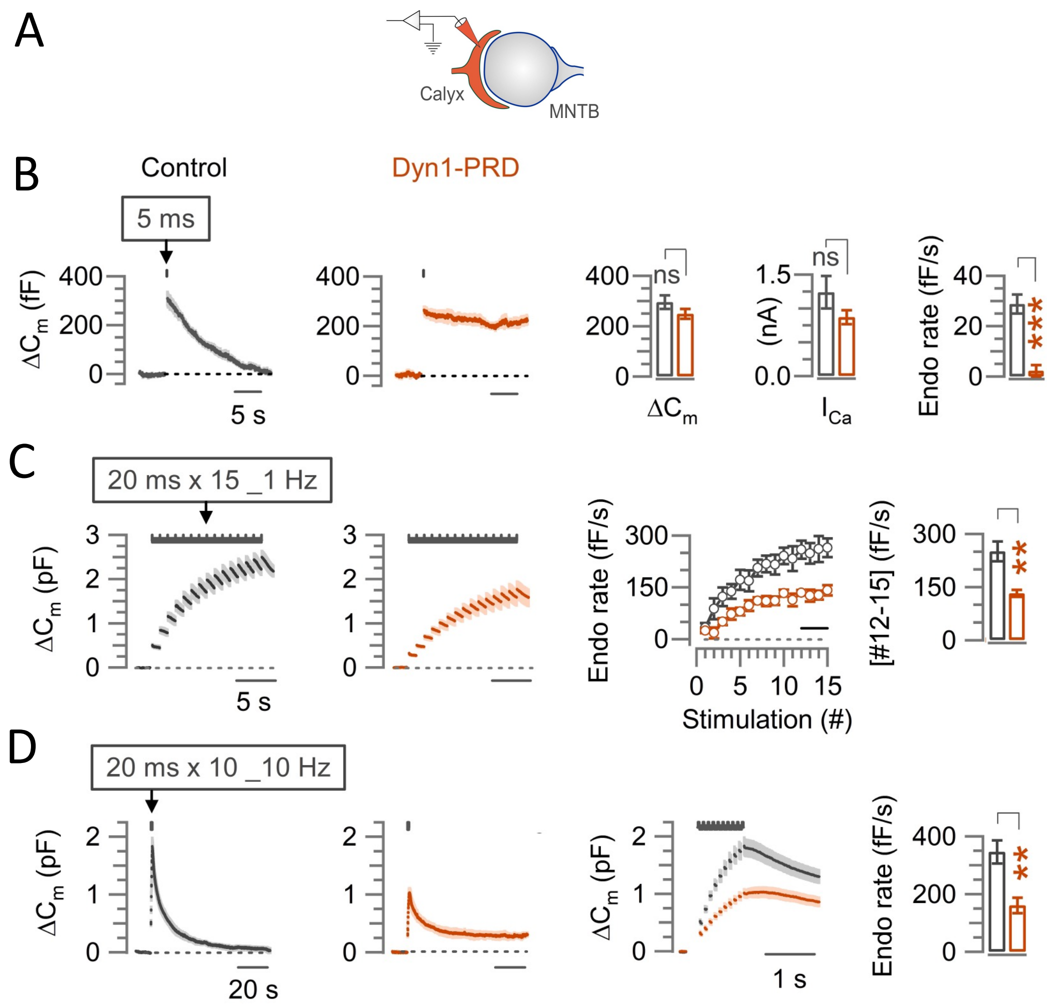
A
Calyx
MNTB
B
Control
Dyn1-PRD
5 ms
ΔCm (fF)
400
200
0
5 s
ns
ΔCm
(nA)
1.5
0.0
ICa
Endo rate (fF/s)
40
20
***
C
20 ms x 15 _1 Hz
ΔCm (pF)
3
2
1
Endo rate (fF/s)
300
150
Stimulation (#)
[#12-15] (fF/s)
**
D
20 ms x 10 _10 Hz
20 s
1 s
400
200

### Figure S2

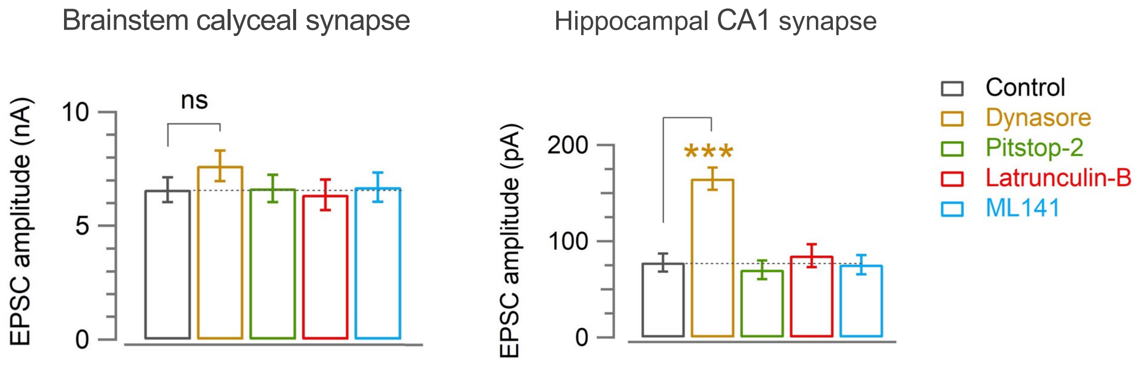
Brainstem calyceal synapse
Hippocampal CA1 synapse
EPSC amplitude (nA)
10
5
0
ns
EPSC amplitude (pA)
200
100
0
***
Control
Dynasore
Pitstop-2
Latrunculin-B
ML141

### Figure S3

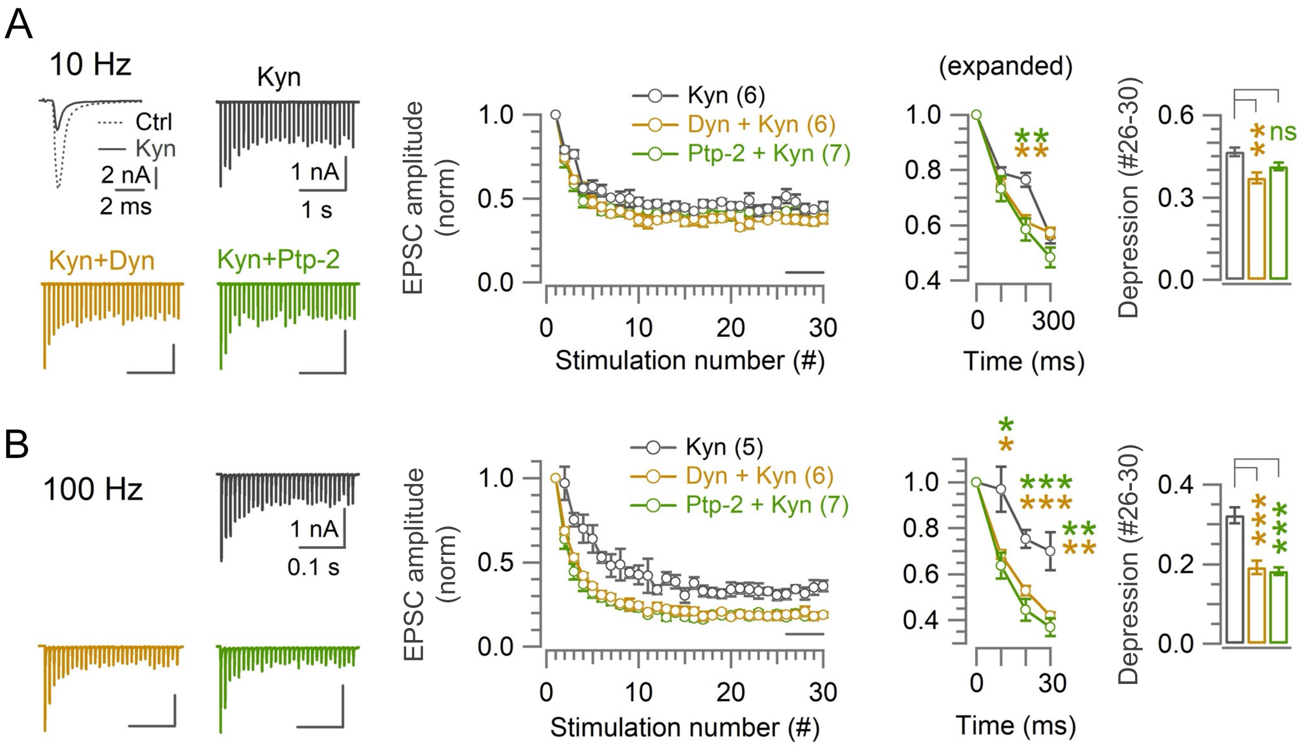

A
10 Hz
Ctrl
Kyn
2 nA
2 ms
Kyn
1 nA
1 s
Kyn+Dyn
Kyn+Ptp-2
Kyn (6)
Dyn + Kyn (6)
Ptp-2 + Kyn (7)
EPSC amplitude (norm)
Stimulation number (#)
(expanded)
Time (ms)
Depression (#26-30)
ns
B
100 Hz
Kyn
1 nA
0.1 s
Kyn (5)
Dyn + Kyn (6)
Ptp-2 + Kyn (7)
EPSC amplitude (norm)
Stimulation number (#)
Time (ms)
Depression (#26-30)

### Figure S4

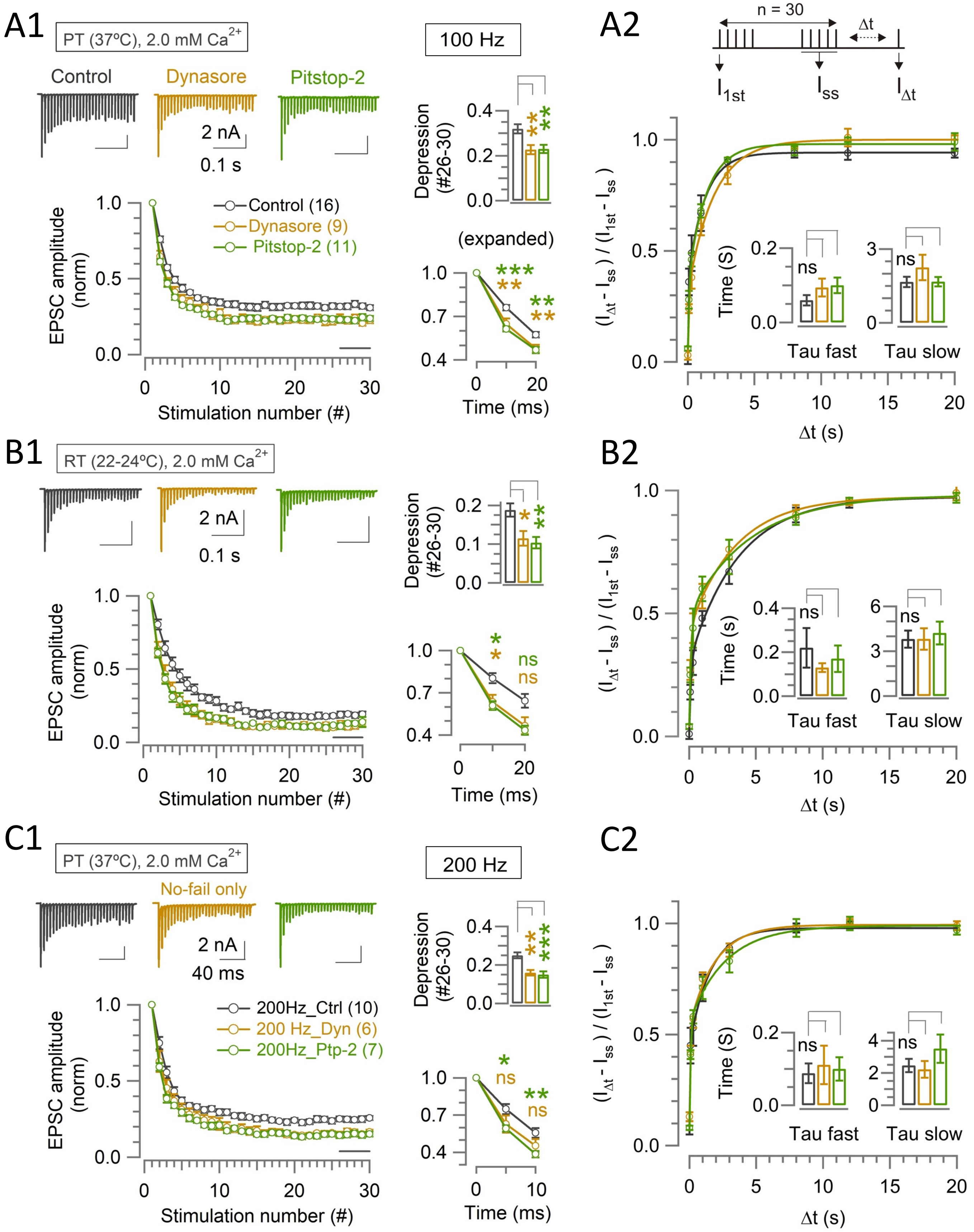

A1
PT (37ºC), 2.0 mM Ca2+
Control
Dynasore
Pitstop-2
2 nA
0.1 s
100 Hz
Depression (#26-30)
(expanded)
EPSC amplitude (norm)
Control (16)
Dynasore (9)
Pitstop-2 (11)
Stimulation number (#)
Time (ms)
A2
n = 30
Δt
I1st
Iss
IΔt
(IΔt - Iss) / (I1st - Iss)
Time (S)
ns
Tau fast
Tau slow
Δt (s)
B1
RT (22-24ºC), 2.0 mM Ca2+
2 nA
0.1 s
Depression (#26-30)
EPSC amplitude (norm)
Stimulation number (#)
ns
Time (ms)
B2
(IΔt - Iss) / (I1st - Iss)
Time (s)
ns
Tau fast
Tau slow
Δt (s)
C1
PT (37ºC), 2.0 mM Ca2+
No-fail only
2 nA
40 ms
200 Hz
Depression (#26-30)
EPSC amplitude (norm)
200Hz_Ctrl (10)
200 Hz_Dyn (6)
200Hz_Ptp-2 (7)
Stimulation number (#)
ns
Time (ms)
C2
(IΔt - Iss) / (I1st - Iss)
Time (S)
ns
Tau fast
Tau slow
Δt (s)

### Figure S5

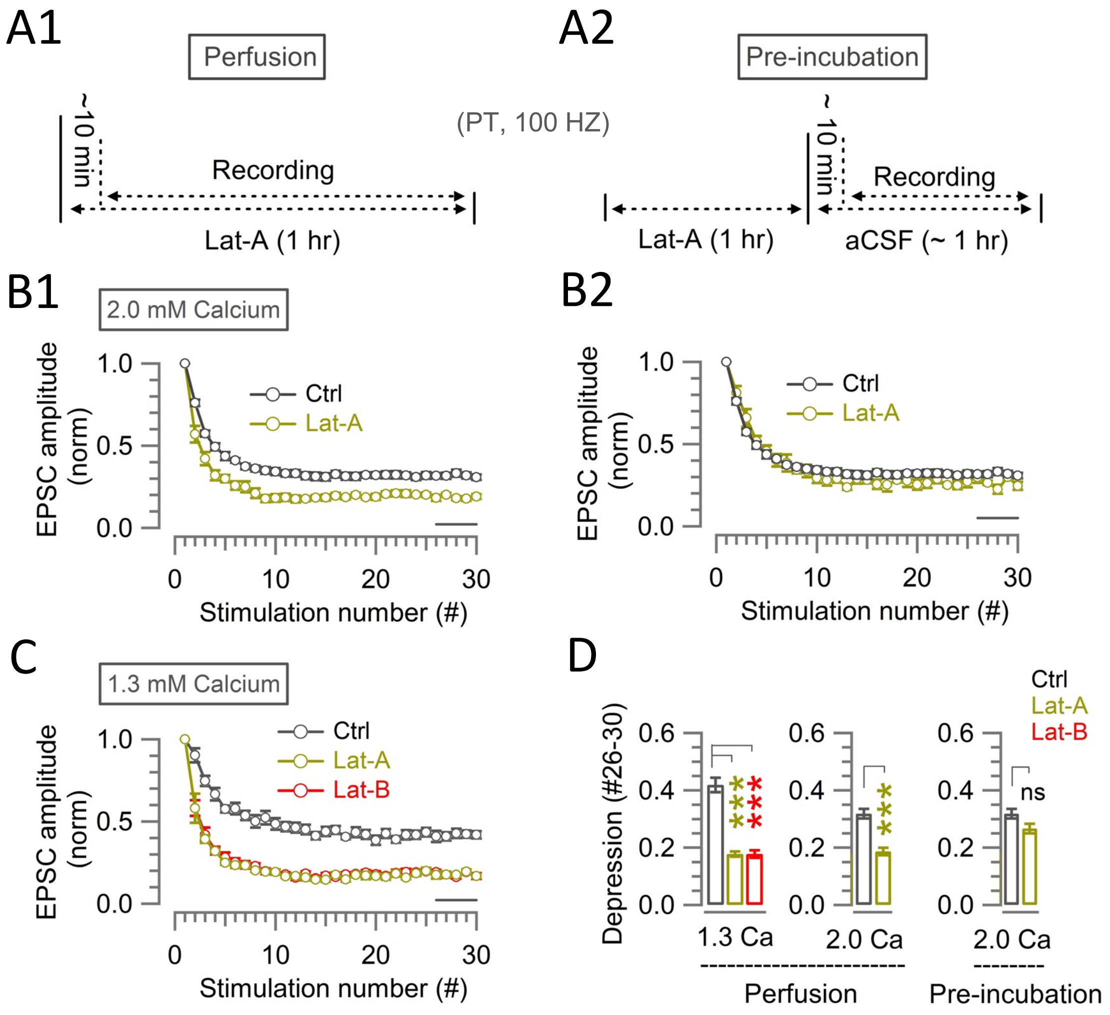
A1
Perfusion
~10 min
Recording
Lat-A (1 hr)
(PT, 100 HZ)
A2
Pre-incubation
~ 10 min
Recording
Lat-A (1 hr)
aCSF (~ 1 hr)
B1
2.0 mM Calcium
EPSC amplitude (norm)
Ctrl
Lat-A
Stimulation number (#)
B2
EPSC amplitude (norm)
Ctrl
Lat-A
Stimulation number (#)
C
1.3 mM Calcium
EPSC amplitude (norm)
Ctrl
Lat-A
Lat-B
Stimulation number (#)
D
Ctrl
Lat-A
Lat-B
Depression (#26-30)
ns
1.3 Ca
2.0 Ca
2.0 Ca
Perfusion
Pre-incubation

### Figure S6

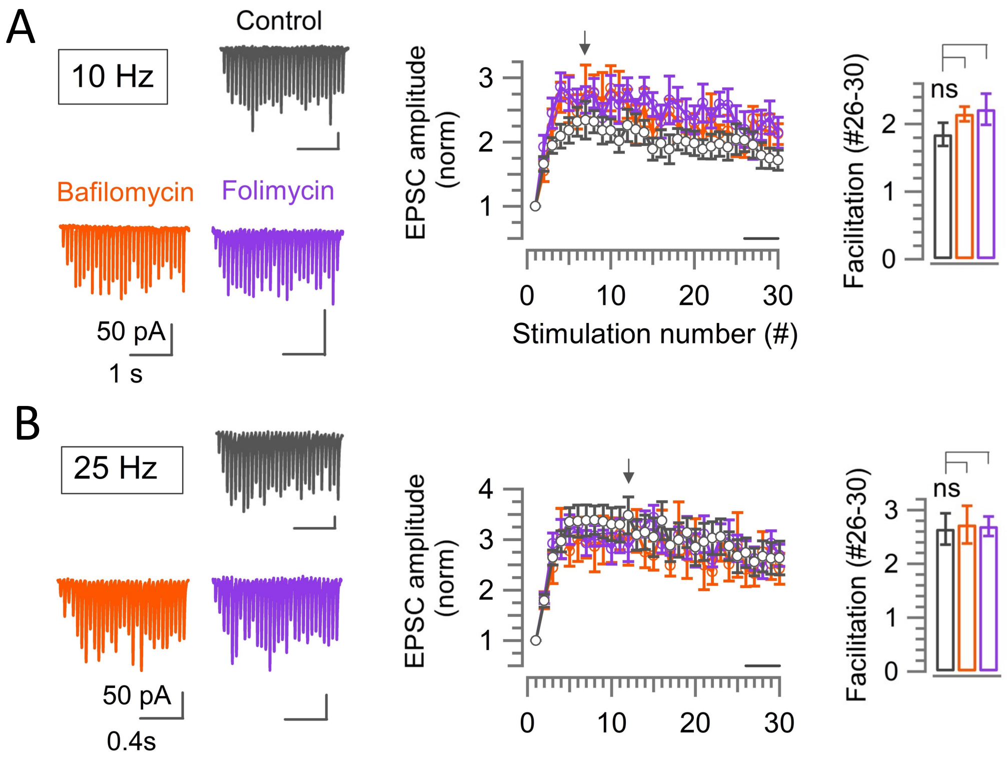
A
10 Hz
Control
Bafilomycin
Folimycin
50 pA
1 s
EPSC amplitude (norm)
Stimulation number (#)
0
10
20
30
1
2
3
Facilitation (#26-30)
ns
0
1
2
B
25 Hz
50 pA
0.4s
EPSC amplitude (norm)
1
2
3
4
0
10
20
30
Facilitation (#26-30)
ns
0
1
2
3

### Table S1

| Parameters | | | Control | Dynasore | Pitstop-2 | Dyn-1 PRD | ML141 | Lat-B |
|---|---|---|---|---|---|---|---|---|
| 5 ms | ΔCm (fF) | Mean (± SEM) | 295 (± 27) | 280 (± 32) | 236 (± 30) | 249 (± 19) | 260 (± 47) | 343 (± 45) |
| | | *p* -value | - | 0.73 | 0.18 | 0.20 | 0.54 | 0.40 |
| | ICa (nA) | Mean (± SEM) | 1.24 (± 0.24) | 1.05 (± 0.13) | 1.05 (± 0.21) | 0.87 ± (0.11) | 0.88 (± 0.29) | 0.98 (± 0.2) |
| | | *p*- value | - | 0.51 | 0.58 | 0.21 | 0.36 | 0.43 |
| | Endo rate (fF/s) | Mean (± SEM) | 28.8 (± 3.7) | 10.3 (± 2.8) | 10.1 (± 3.4) | 2.3 (± 2.2) | 23.0 (± 5.8) | 25.6 (± 5.2) |
| | | *p* -value | - | 0.004 | 0.006 | < 0.001 | 0.44 | 0.65 |
| | | (n) | 5 | 5 | 5 | 5 | 6 | 6 |
| 20 ms x 15_1 Hz | Cumulative ΔCm (pF) | Mean (± SEM) | 2.41 (± 0.16) | 1.25 (± 0.08) | 1.67 (± 0.08) | 1.7 (± 0.21) | 1.87 (± 0.1) | 2.05 (± 0.3) |
| | | *p* -value | - | 0.0007 | 0.004 | 0.02 | 0.02 | 0.28 |
| | Endo rate (fF/s) | Mean (± SEM) | 251 (± 28.3) | 121 (± 19.1) | 133 (± 15.6) | 132 (± 9.8) | 214 (± 19.1) | 233 (± 47.6) |
| | (Last four points average) | *p* -value | - | 0.004 | 0.007 | 0.001 | 0.3 | 0.7 |
| | | (n) | 6 | 6 | 5 | 7 | 5 | 4 |
| 20 ms x 10_10 Hz | Cumulative ΔCm (pF) | Mean (± SEM) | 1.67 (± 0.15) | 1.03 (± 0.09) | 1.0 (± 0.12) | 1.03 (± 0.1) | 1.31 (± 0.11) | 1.42 (± 0.3) |
| | | *p* -value | - | 0.008 | 0.009 | 0.007 | 0.09 | 0.49 |
| | Endo rate (fF/s) | Mean (± SEM) | 346 (± 40.1) | 204 (± 25.3) | 131 (± 24.6) | 161 (± 27.3) | 242 (± 21.5) | 350 (± 33.3) |
| | | *p* -value | - | 0.017 | 0.004 | 0.003 | 0.08 | 0.95 |
| | | (n) | 6 | 4 | 4 | 6 | 4 | 5 |
| Figure(s): | | | 1, 3 and S1 | 1 | | S1 | 3 | |

Endocytosis rate was measured from following time frame(s)

1) 5 ms: (0.45 - 5.45) s after stimulation
2) 20 ms x 15_1 Hz: (0.45 - 0.95) s after every stimulation
3) 20 ms x 10_10 Hz: (0.45 - 1.45) s after last stimulation

### Table S2

| Frequency | Conditions/parameters | | | Tau fast (s) | Tau slow (s) | Tau mean (s) | Af/As | (n) | Figure (s) |
|---|---|---|---|---|---|---|---|---|---|
| 10 Hz | IΔt/I1st | Control | Mean (± SEM) | - | - | 2.3 (± 0.4) | - | 9 | |
| | | Dynasore | Mean (± SEM) | - | - | 1.7 (± 0.2) | - | 8 | |
| | | | *p* -value | | | 0.2 | | | |
| | | Pitstop-2 | Mean (± SEM) | - | - | 1.9 (± 0.3) | - | 6 | |
| | | | *p* -value | | | 0.5 | | | |
| | | ML141 | Mean (± SEM) | - | - | 2.6 (± 0.2) | - | 8 | |
| | | | *p* -value | | | 0.61 | | | |
| | | Lat-B | Mean (± SEM) | - | - | 3.7 (± 0.9) | - | 6 | |
| | | | *p* -value | | | 0.14 | | | |
| | (IΔt - Iss)/(I1st - Iss) Normalized at Iss | Control | Mean (± SEM) | - | - | 2.2 (± 0.4) | - | 9 | 2,4 |
| | | Dynasore | Mean (± SEM) | - | - | 1.8 (± 0.3) | - | 8 | 2 |
| | | | *p* -value | | | 0.32 | | | |
| | | Pitstop-2 | Mean (± SEM) | - | - | 2.1 (± 0.24) | - | 6 | 2 |
| | | | *p* -value | | | 0.9 | | | |
| | | ML141 | Mean (± SEM) | - | - | 3.0 (± 0.5) | - | 8 | 4 |
| | | | *p* -value | | | 0.21 | | | |
| | | Lat-B | Mean (± SEM) | - | - | 3.4 (± 0.6) | - | 6 | 4 |
| | | | *p* -value | | | 0.16 | | | |
| 100 Hz | IΔt/I1st | Control | Mean (± SEM) | 0.21 (± 0.06) | 3.5 (± 0.5) | 1.9 (± 0.4) | 1.4 (± 0.3) | 8 | |
| | | Dynasore | Mean (± SEM) | 0.069 (± 0.01) | 2.0 (± 0.3) | 0.93 (± 0.06) | 1.3 (± 0.2) | 9 | |
| | | | *p* -value | 0.02 | 0.008 | 0.01 | 0.7 | | |
| | | Pitstop-2 | Mean (± SEM) | 0.06 (± 0.014) | 1.8 (± 0.24) | 0.8 (± 0.1) | 1.5 (± 0.4) | 6 | |
| | | | *p* -value | 0.034 | 0.008 | 0.02 | 0.8 | | |
| | | ML141 | Mean (± SEM) | 0.14 (± 0.03) | 2.3 (± 0.2) | 1.3 (± 0.2) | 1.1 (± 0.2) | 8 | |
| | | | *p* -value | 0.23 | 0.04 | 0.17 | 0.45 | | |
| | | Lat-B | Mean (± SEM) | 0.07 (± 0.014) | 2.6 (± 0.5) | 1.43 (± 0.2) | 0.9 (± 0.15) | 8 | |
| | | | *p* -value | 0.04 | 0.2 | 0.3 | 0.2 | | |
| | (IΔt - Iss)/(I1st - Iss) Normalized at Iss | Control | Mean (± SEM) | 0.18 (± 0.05) | 3.2 (± 0.5) | 1.8 (± 0.4) | 1.5 (± 0.4) | 8 | 2,4 |
| | | Dynasore | Mean (± SEM) | 0.05 (± 0.009) | 1.5 (± 0.2) | 0.78 (± 0.09) | 0.9 (± 0.12) | 9 | 2 |
| | | | *p* -value | 0.008 | 0.003 | 0.02 | 0.24 | | |
| | | Pitstop-2 | Mean (± SEM) | 0.04 (± 0.014) | 1.3 (± 0.3) | 0.53 (± 0.08) | 1.6 (± 0.4) | 6 | 2 |
| | | | *p* -value | 0.02 | 0.008 | 0.02 | 0.8 | | |
| | | ML141 | Mean (± SEM) | 0.13 (± 0.024) | 2.3 (± 0.3) | 1.3 (± 0.2) | 1.1 (± 0.2) | 8 | 4 |
| | | | *p* -value | 0.4 | 0.15 | 0.2 | 0.4 | | |
| | | Lat-B | Mean (± SEM) | 0.07 (± 0.02) | 2.5 (± 0.4) | 1.6 (± 0.3) | 0.73 (± 0.23) | 8 | 4 |
| | | | *p* -value | 0.05 | 0.4 | 0.63 | 0.42 | | |

Time constants were obtained by fitting recovery points of individual cells and taking average across cells.

| | | number of cells (n) at Δt | | | | | | | |
|---|---|---|---|---|---|---|---|---|---|
| Frequency | Δt (s) -> | 0.02 | 0.1 | 0.3 | 1 | 3 | 8 | 12 | 20 |
| 10 Hz | Control | - | - | 11 | 10 | 10 | 10 | 9 | 8 |
| | Dynasore | - | - | 8 | 8 | 8 | 8 | 8 | 6 |
| | Pitstop-2 | - | - | 7 | 7 | 7 | 7 | 7 | 7 |
| | ML141 | - | - | 9 | 8 | 8 | 8 | 8 | 8 |
| | Lat-B | - | - | 7 | 7 | 7 | 7 | 7 | 7 |
| 100 Hz | Control | 11 | 10 | 10 | 9 | 10 | 9 | 5 | 4 |
| | Dynasore | 10 | 10 | 9 | 9 | 9 | 9 | 7 | 7 |
| | Pitstop-2 | 6 | 7 | 7 | 7 | 7 | 7 | 7 | 7 |
| | ML141 | 10 | 8 | 8 | 8 | 8 | 8 | 8 | 8 |
| | Lat-B | 8 | 8 | 8 | 8 | 8 | 8 | 8 | 8 |

Recovery curves in Fig 2,4, and S4, were obtained by fitting the average recovery points at every Δt time.
Individual cells that recovered completely within 8 s or 12 s, without having further data points (at 12 s and/or 20 s), were extrapolated for fitting.
